# Serotonin signaling in hippocampus during initial cocaine abstinence drives persistent drug seeking

**DOI:** 10.1101/279729

**Authors:** Amy S. Kohtz, Gary Aston-Jones

## Abstract

**Background:** The initiation of abstinence (extinction day 1, ED1) represents a stressful event involving abstinence from drug. We showed previously that ED1 cocaine-seeking behavior is reduced by blocking 5-HT signaling in dorsal hippocampus in both male and female rats. We hypothesized that the experience of ED1 can substantially influence later relapse behavior, and that dorsal raphe serotonin (DR 5-HT) signaling to dorsal hippocampus (DH) may be involved.

**Methods:** We used pharmacological inhibition of dorsal hippocampus 5-HT1A/1B receptors (via WAY100,635 plus GR127935), and chemogenetic inhibition of dorsal raphe-dHPC signaling to test the roles of these pathways on cocaine-seeking 2 weeks after ED1. We also tested specific effects of 5-HT1A or 5-HT1B receptor antagonism on conditioned place preference for cocaine.

**Results:** Inhibition of DR-DH signaling via DREADDs or 5-HT1A/1B antagonists decreased ED1 drug-seeking with persistent effects on cocaine-seeking 2 weeks later, confirming the involvement of 5-HT signaling to dorsal hippocampus in driving drug-seeking persistence. Administration of a 5-HT1B antagonist alone on ED1 transiently decreased drug-associated memory performance in CPP, whereas administration of a 5-HT1A antagonist had no effect on memory but blocked CPP on a subsequent test 24h later.

**Conclusions:** We conclude that blockade of DR inputs or 5HT1 signaling in DH on ED1 prevents recall of the drug-associated context and reduces drug seeking via antagonism of 5-HT1B receptors, and consolidates the memory of the newly non-drug context via antagonism of 5-HT1A receptors. Thus, treatments that modulate 5-HT-dependent memory mechanisms during initial abstinence may facilitate later maintenance of abstinence.

## Introduction

Cocaine withdrawal-related anxiety is highest during the first 2 days of cessation of use (Coffey et al., 2000), and craving during this initial abstinence period can predict later relapse propensity (Hall et al., 1991). Therefore, initial drug abstinence may be a critical time point for understanding factors that contribute to later relapse.

Prior reports implicate corticotrophin-releasing factor-1 (CRF-1), norepinephrine (NE), and serotonin signaling (5-HT) in driving cocaine-seeking behavior during initial abstinence as seen on extinction day 1 (ED1) (Cason et al., 2016, Kohtz and Aston-Jones, 2017). These studies also show that cocaine-seeking on ED1 is marked by increased Fos expression in locus coeruleus (LC), dorsal raphe (DR), median raphe, and dorsal hippocampus (DH). In addition, antagonism of β-adrenergic receptors in DH decreased ED1 cocaine-seeking in females only, whereas blocking 5-HT1 receptors in DH on ED1 substantially decreased ED1 drug-seeking in both sexes (Kohtz and Aston-Jones, 2017). However, the effects of manipulating these circuits during ED1 on later cocaine seeking remains unknown.

Our study employs a novel approach that focuses not only on cocaine-seeking during ED1, but also on the persistence of drug seeking following 2 weeks of abstinence. Our seeking-persistence paradigm (SPP) can be used to study either transient (during ED1) or persistent effects (longer abstinence) of manipulating ED1 on cocaine seeking behaviors. We focused on serotonin signaling in the present study, given prior reports implying its role in both sexes to modulate cocaine-seeking.

## Methods

### Jugular Catheter Surgeries

Animals were anesthetized with ketamine/xylazine (56.5/8.7 mg/kg) and given rimadyl (1 mg/kg) as an analgesic. Chronic in-dwelling catheters were constructed in-house and inserted as described previously (Smith et al, 2009). Animals were given cefazolin (10 mg, intravenous (i.v.)) and heparin (10 U, i.v.) starting 3 days following surgery and daily following self-administration sessions. Rats recovered from surgery for at least 1 week before self-administration training.

### Dorsal Hippocampus Cannulae and Virus Injections

Immediately following jugular catheter surgery rats were placed in a stereotaxic apparatus and bilaterally infused via glass micropipette with 1uL per side of AAV8-Syn-hM4Di-mCherry targeted to the dorsal raphe (male coordinates: AP: -7.8, ML: ±3.1, DV: -7.5, 30° angle; female coordinates: AP: -7.6, ML: ±3.1, DV: -7.2, 20° angle). Immediately following virus implantation, bilateral guide cannulae were implanted targeted to DH (AP: -3.0, ML: ±2.0, DV:-2.5, no angle). Cannula placements and DREADD expression were confirmed for each rat following completion of behavioral analysis (details below).

### Drugs

Cocaine hydrochloride (NIDA, Research Triangle Park, NC) was dissolved in 0.9% sterile saline. WAY100,635 (Tocris) and GR127935 (Tocris) were dissolved in 0.9% sterile saline and administered intrahippocampally (IHC; 1 nmol/1.0 µl/side) at a rate of 0.25 µl/min, 10 min before testing (Kohtz and Aston-Jones, 2017). CNO (supplied by RTI International, North Carolina) was dissolved in sterile aCSF and 10mM was administered in a volume of 1uL per side.

### Cocaine self-administration

Self-administration sessions were conducted in standard operant chambers housed in sound attenuating cubicles and controlled via MED-PC IV software (Med-Associates, St Albans, VT) as described previously (Smith and Aston-Jones, 2011). Rats were trained in daily 2hr sessions to press an active lever for intravenous cocaine (0.2 mg/50 ul infusion for males, 0.16 mg/50 ul infusion for females) for at least 10 days on a fixed ratio 1 (FR1) schedule of reinforcement to reach a criterion of >10 cocaine infusions/day. Each cocaine infusion was followed by a 20 second time-out period in which lever pressing produced neither cocaine nor cues. An inactive lever was also present; presses on it were tabulated but had no consequence.

### Cocaine-seeking during ED1

Twenty-four hours after the final self-administration session, rats were tested in an initial extinction session (ED1). In this session rats were exposed to the self-administration chamber for 90 min during which time presses on either lever had no consequence. Active lever pressing in the absence of cocaine reward or cues on ED1 was taken as a measure of cocaine-seeking during early abstinence, as previously reported (Feltenstein et al., 2011, Cason et al., 2016, Kohtz and Aston-Jones, 2017).

### Cocaine Seeking Persistence

Following the ED1 test, rats were returned to the homecage for 14 days of abstinence. Then, rats were returned to the context as in ED1 procedures, for a second extinction session (ED2) on withdrawal day 15 (WD15). Active lever pressing during this second session was taken as a measure of cocaine seeking persistence following abstinence.

### Conditioned Place Preference (CPP)

We used a standard two-chamber balanced design. Rats were habituated and assessed for a side preference prior to conditioning. The cocaine-paired chamber was counterbalanced between rats. Two conditioning sessions (for saline and cocaine) occurred for 3 days consecutively, prior to testing. Injections of cocaine or saline alternated between morning and afternoon sessions. Conditioning and testing were identical to our previously published reports (Harris and Aston-Jones, 2003b, a).

### Immunohistochemistry

Immunohistochemistry was used to confirm localization of the AAV8-Synapsin-Hm4Di-mCherry receptor expression in cell bodies of the dorsal raphe and processes to dorsal hippocampus. Co-staining for mCherry and 5-HT was performed in the dorsal raphe to confirm containment of transduction sites. As a synapsin promoter was used, some expression was observed outside the borders of the dorsal raphe in most animals. Animals with more than ∼30% of DREADD expression observed outside dorsal raphe were excluded from analyses (n=2). Misses are not reported in the current manuscript, as only 2 rats with misses survived surgical recovery and operant testing. Loss of rats within 1 week of surgery amounted to 6 rats that we attributed to puncture of the aqueduct or aqueduct pressure, given the proximity of the dorsal raphe to the aqueduct. In included rats, hM4Di expression was robust in dorsal raphe (Fig.2A). Immunohistochemistry for mCherry (Mahler et al., 2014) and 5-HT (Kohtz and Aston-Jones, 2017) were performed per previously reported methods.

### Statistical Analysis

Independent t-tests, Pearson,s R correlations, one- or two-way analysis of variance (ANOVA), or three-way multiple ANOVAs (MANOVA) were used to compare differences between groups in responding.

## Results

### Sex-specific cocaine infusion doses (0.2mg/infusion for males, 0.16mg/infusion for females) produced similar self-administration behavior in male and female rats

We previously found that 0.2mg/infusion in male rats and 0.16mg/infusion in female rats produced statistically similar average lever pressing behavior (∽50 infusions/2 hr session) during the last 3 days of self-administration (Kohtz and Aston-Jones, 2017). Our prior report found that average lever pressing in females was greater during the first 3 days of self-administration compared to males (Kohtz and Aston-Jones, 2017). Similarly, here we found that females had more overall lever pressing behavior (main effect of sex: F(1,26)=4.303, *p* < 0.05); post-hoc *t-tests* revealed that this difference was only significant on day 5 (post-hoc t=3.710, *p* < 0.05) (Fig.1B). There was also a significant interaction between sex and self-administration day for inactive lever pressing (F(1,26)=2.024, *p* < 0.05), wherein females had greater early inactive lever pressing than males. Post-hoc *t-tests* indicated that males and females significantly differed in inactive lever pressing on day 4 only (post-hoc t=3.710, *p* < 0.05) (Fig.1B). There was a main effect of self-administration day on number of infusions of cocaine obtained, in that number of infusions increased over time in both sexes (F(1,26)=6.490, *p* <0.05) (Fig.1B). We also show that females had more overall cocaine self-administration (mg/kg/day; main effect of sex: F(1,26)=10.29, *p* < 0.05); post-hoc *t-tests* revealed that this difference was only significant on days 4-7 (post-hoc t=3.051, t=3.978, t=3.563, t=3.210, *p* < 0.05) driving a significant interaction between sex and day (F(1,26)=2.443, *p* < 0.05; Fig.1C).

### Persistent Effects of Pharmacological Antagonism of 5-HT1A and 5-HT1B Receptors in the Dorsal Hippocampus on Cocaine-Seeking

Following 10 days of >10 infusions per day during cocaine self-administration, rats (n=6-9/group) were returned to the operant chamber without cues or cocaine reward for ED1 testing. 10 minutes prior to entering the operant chamber, rats were given an intra-hippocampal infusion of saline or a cocktail of WAY100,635 plus GR127935 (5-HT1A and 5-HT1B antagonists, respectively, WAY/GR; 1 nmol each/1.0 µl/side). Our prior report indicated that such infusions reduced active lever pressing without locomotor effects (Kohtz and Aston-Jones, 2017). As previously reported, there was a significant main effect of sex wherein females had greater ED1 responding than males (F(1,24)=6.492, *p* <0.05) (Kohtz and Aston-Jones, 2017). There was also a significant main effect of treatment, wherein administration of WAY/GR decreased cocaine-seeking behavior on ED1 in both males and females (F(1,24)=28.09, *p* <0.05) (Fig.1D). Rats were then returned to the homecage for 2 weeks of forced abstinence. On withdrawal day 15, rats were returned to the operant chamber for a second time without drugs or cues (ED2) to test cocaine-seeking persistence, with no further hippocampal or other injections. To test the effects of ED1 WAY/GR on cocaine-seeking persistence, we ran a three-way repeated measures MANOVA comparing sex and treatment over ED1 and ED2 test days. There was a significant main effect of sex (F(1,48)=5.800, *p* < 0.05) as well as a main effect of treatment (F(1,48)=48.650, *p* <0.05), but no main effect of test day (F(1,48)=5.216, NS) (Fig.1D). There was no significant interaction between sex and treatment across ED1 and ED2 (F(1,48)=1.473, NS). These data indicate that females have greater cocaine-seeking than males and intervention with WAY/GR on ED1 decreases cocaine-seeking behavior on both ED1 and ED2, showing that attenuation of 5HT signaling on ED1 persistently decreased drug seeking.

Separate groups of rats (n=11-12/sex) were trained in self-administration for 10 days of >10 infusions/day as described above. Twenty-four hr following the final operant session, rats were infused with WAY/GR in DH and returned to their homecage to test if exposure to the context on ED1 was required for the effects of the 5HT antagonists on persistent cocaine-seeking. Rats then underwent 14 days of abstinence, and were tested in a delayed ED1 (on WD15) with no further manipulations. Active lever presses were recorded as a measure of cocaine-seeking behavior. There was no significant interaction between sex and treatment (F(1,19)=0.0604, NS), nor significant main effects of sex (F(1,19)=0.8609, NS) or treatment (F(1,19)=1.666, NS) on lever pressing (Fig.1F). This indicates that exposure to the context is necessary for WAY/GR treatment to reduce persistent cocaine-seeking, tested 2wks later.

As females are reported to be more extinction resistant than are males (Lynch and Carroll, 2000, Becker and Hu, 2008), we investigated whether infusion with WAY/GR on ED1 altered sex differences in extinction resistance. Initial extinction burst responding on ED1 is greater in females compared to males, however cocaine-seeking on ED2 is similar in both under saline conditions (t=5.391, *p* <0.05; Fig.1D). As described above, both initial cocaine-seeking on ED1 and on ED2 were decreased by WAY/GR administration. Following ED2 testing (Fig.1D), all rats were extinguished in the operant box for at least 7 days (no cues) until the criterion of <25 lever presses (active+inactive lever presses) was met for 3 consecutive days. Survival analyses revealed an effect of sex and treatment (saline or WAY/GR on ED1) (X^2^=11.45, *p* < 0.05) wherein WAY/GR in DH on ED1 decreased extinction resistance in females but not in males (Fig.1G). Post-hoc tests in females reveal an effect of treatment (X^2^=12.96, *p* < 0.05) on extinction resistance, wherein females treated with saline took longer to meet extinction criterion than did males.

Rats that were extinguished (i.e., Fig.1G) were tested for cocaine-seeking during cued reinstatement. First, rats were administered saline 10 mins prior to being placed in the operant chamber for a 2 h cued reinstatement session, wherein active lever presses yielded cocaine-associated cues but no cocaine. There were no significant effects of the prior ED1 treatment (e.g. saline vs WAY/GR) on cued reinstatement responding in males or females (F(1,23)=1.673, NS; Fig.1H).

Rats were subjected to additional extinction training (criterion of <25 lever presses (active+inactive lever presses) for 3 consecutive days) and then tested for cued reinstatement responding 10 min following WAY/GR infusion into DH. A two-way repeated measures ANOVA comparing the effects of sex and acute treatment found an overall effect of sex, wherein females had greater cued-reinstatement behavior than did males (F(1,50)=4.571, *p* < 0.05), but no effect of WAY/GR treatment on reinstatement test day on cued reinstatement behavior (F(1,50)=0.0214, NS; Fig.1H).

### Persistent Effects of chemogenetic inhibition of dorsal raphe input to Dorsal Hippocampus on Cocaine Seeking

Twenty-two rats were infused with an AAV8-hSyn-hM4D(Gi)-mCherry in dorsal raphe as described in Methods. Two rats were excluded from analyses due to substantial DREADD expression outside of dorsal raphe, and 6 rats did not survive surgery. Rats were also implanted with cannulae in DH and intravenous catheters at the time of virus surgery. Rats began self-administration training 4-5 weeks following surgery, to allow time for DREADD expression and transport. As above, rats were trained until they achieved 10 days of >10 cocaine infusions per day during 2h self-administration sessions. The following day, rats (n=6-8/group) were returned to the operant chamber for a 2h ED1 test session without cues or cocaine reward. Active and inactive levers were extended, and presses on the active lever were used to indicate cocaine-seeking behavior. Ten minutes prior to entering the operant chamber, rats were given an intra-hippocampal infusion of aCSF (1.0uL/side) or CNO (1mM, 1.0uL/side), to test the effects of inhibiting dorsal raphe signals to DH. As in the 5-HT antagonist studies described above, there was a main effect of sex, wherein females had greater ED1 responding than males (F(1,24)=26.28, *p* <0.05), and a significant main effect of treatment, wherein administration of CNO decreased cocaine-seeking behavior on ED1 in both males and females (F(1,24)=12.88, *p* <0.05) (Fig.2E). There also was a significant interaction between sex and treatment condition, wherein intra-DH CNO was more effective in females than in males (F(1,24)=9.420, *p* <0.05).

Following this ED1 test, rats were returned to their homecage for 2 weeks of forced abstinence. On withdrawal day 15, rats were returned to the operant chamber for a second extinction test (ED2) to test cocaine-seeking persistence, with no further injections. We used a three-way repeated measures MANOVA to compare sex and treatment over the ED1 and ED2 test days. We found a significant main effect of sex (F(1,48)=16.856, *p* < 0.05) as well as a main effect of treatment (F(1,48)=53.716, *p* <0.05) across ED1 and ED2, but no main effect of test day (F(1,48)=0.182, NS). We also found a significant interaction between sex and treatment across ED1 and ED2 (F(1,48)=8.482, *p* <0.05). Similar to our pharmacology manipulations, we found no effect of CNO to influence cued-reinstatement behavior, however did find a main effect of sex wherein females had greater cued reinstatement than did males (F(1,18)=14.00, *p* <0.05; Fig.2F). These data indicate that inhibition of the endogenous dorsal raphe signal to dorsal hippocampus reduces cocaine seeking persistently in males and, to a greater extent, in females.

### 5-HT1A and 5-HT1B Receptor Antagonists in dorsal hippocampus decrease recall and consolidation of drug context memory: Conditioned Place Preference experiments

In the seeking persistence paradigm, blockade of dorsal raphe signaling in dorsal hippocampus CA1 via DREADDs or 5-HT1A/1B antagonists reduced cocaine seeking persistently. We speculated that the reduction in cocaine seeking on ED1 by these treatments reflected reduced recall of the drug-associated context. We also hypothesized that these treatments blocked the consolidation of new information on ED1 (e.g. that context was no longer associated with drug) because there was a persistent decrease in drug seeking 2 weeks later. We investigated these hypotheses by testing the effects of intra-hippocampal 5-HT antagonists given together, or separately, in the conditioned place preference paradigm (CPP).

A naïve group of rats was used to test the effects of 5-HT1A or 5-HT1B receptor antagonism in dorsal hippocampus on CPP for cocaine. There was a main effect of treatment when the WAY/GR cocktail was administered 10 minutes prior to each CPP conditioning session, wherein CPP on test day was enhanced (F(1,23)=11.72 *p* <0.05) in both sexes (Fig.3B). In contrast, when WAY/GR was administered 10 minutes prior to the CPP test session only, there was a main effect to reduce cocaine CPP (F(1,24)=21.77, *p* <0.05) similarly in both sexes (Fig.3B). These data show that WAY/GR enhanced the acquisition cocaine CPP when given during conditioning, but produced deficits in the expression of cocaine CPP when administered on test day.

As effects were similar in males and females, we investigated the effects of the individual compounds during acquisition vs. expression in naive males only. We found an effect of WAY100,635 (selective 5-HT1A antagonist) to produce time of administration/time of testing dependent enhancements or deficits in CPP (F(3,23)=14.08, *p* <0.05; Fig.3C). WAY administered during CPP acquisition significantly increased cocaine CPP (t=3.321, *df*=10, *p* <0.05), whereas WAY administered the test day had no effect compared to saline administered controls (t=0.6406, *df*=11, NS). When retested 24 hours later, rats that had been administered WAY on test day only showed significant deficits in CPP (t=3.3799, *df*=11, *p* <0.05). These results indicate that WAY in hippocampus facilitates learning an association between context and cocaine reward.

Next, we tested the GR127935 (5-HT1B antagonist) using the same paradigm in an additional group of naïve rats. We found an effect of GR to produce time of administration/time of testing dependent deficits in CPP (F(3,25)=12.56, p <0.05; Fig.3D). GR administration during CPP acquisition had no effect on cocaine CPP (t=0.0392, *df*=10, NS), whereas GR given only on the test day decreased cocaine CPP (t=4.666, *df*=12 *p* <0.05). In contrast to effects of WAY, these effects of GR were transient as CPP on the 24hr-retest was not significantly different from saline controls (t=0.9019, *df*=12, NS).

## Discussion

Our study identifies the role of serotonin signaling in dorsal hippocampus on the persistence of cocaine-seeking and on drug-associated memories. First, we established a paradigm to measure cocaine-seeking following ED1 testing. We recapitulated our prior finding that ED1 cocaine-seeking is attenuated by 5-HT1A and 5-HT1B antagonists in dorsal hippocampus (Kohtz and Aston-Jones, 2017). We extended these findings to show that attenuating 5HT/dorsal raphe signaling in dorsal hippocampus on ED1 via pharmacology or chemogenetic inhibition has persistent effects to reduce cocaine-seeking 2 weeks later. We show that co-exposure to 5HT antagonists in the cocaine self-administration context is required to decrease persistent seeking. We also show specific effects of 5-HT1A vs. 5-HT1B receptors on cocaine-associated contextual memory recall vs consolidation, by using the CPP paradigm.

### Sex Differences in 5-HT Signaling

Prior reports indicate that there are sex differences in hippocampal 5-HT signaling. These studies discuss differences in expression patterns, and functional effects on neurogenesis, depression, stress, and mood disorders (Zhang et al., 1999, Martinowich and Lu, 2008, Franceschelli et al., 2015). In particular, sex differences in hippocampus somatodendritic 5-HT1A receptors play a substantial role in regulating stress responses (Haleem and Parveen, 1994, Haleem, 2011). Indeed, systemic administration of a 5-HT1A agonist produces a two-fold decrease of 5-HT release in hippocampus of females compared to males (Haleem et al., 1990), indicating that there is greater 5-HT1A receptor sensitivity in females compared to males. Acute stressors increase 5-HT1A receptor sensitivity, thereby decreasing 5-HT availability; producing coping deficits (Haleem et al., 2007). Conversely, repeated predictable stress decreases 5-HT1A receptor sensitivity, increasing 5-HT availability; producing stress adaptation (Lopez et al., 1998). Thus, raphe-hippocampal serotonin signaling contributes to stress adaptation in a sex-dependent manner, wherein females may be more sensitive to acute stressors, e.g. ED1. We have previously shown that ED1 is a highly stressful event, producing both a >4-fold increase in circulating corticosterone as well as substantial Fos activation in both the dorsal raphe and dorsal hippocampus (Cason et al., 2016, Kohtz and Aston-Jones, 2017). Herein, we show that increased cocaine-seeking among females on ED1 is persistently attenuated to a greater extent by 5-HT1A/1B antagonism compared to males. Thus, increased cocaine-seeking on ED1 may be driven by greater 5-HT1A receptor sensitivity in the dorsal hippocampus, in particular among females.

### Serotonin Exerts Diverse Effects on Memory Processes

The role of 5-HT signaling in learning and memory is controversial such that the release of 5-HT into the hippocampus can exert both positive and negative effects on multiple memory types, due at least in part to the large number of 5-HT receptors and diverse downstream signaling (Hoyer and Martin, 1997, Hoyer et al., 2002). One such controversial result is that both increased 5-HT activity and inhibition of brain 5-HT synthesis can impair learning performance, indicating that 5-HT activity has U-shaped effects on memory (Ogren et al., 1982, Ogren, 1985). Specific roles of 5-HT1A and 5-HT1B receptors in dorsal hippocampus CA1 pyramidal and interneurons on memory acquisition and performance are of particular interest to the interpretation of results found in the current manuscript.

### CA1 5-HT1A Receptors in Memory Acquisition and Consolidation

Activation of CA1 5-HT1A receptors impairs spatial memory acquisition, and can be reversed by administration of 5-HT1A specific antagonists such as WAY100,635 (Carli et al., 1999, Eriksson et al., 2007). Further support for an inhibitory role of 5-HT1A receptors on memory is reviewed in (Ogren et al., 2008). Consistent with this view, we show that the 5-HT1A receptor antagonist WAY100,635 administered during cocaine CPP acquisition (but not immediately prior to testing) enhanced cocaine-associated contextual memory. A 24-hr retest further supported this interpretation, as 5-HT1A antagonism on test day produced deficits in CPP 24hrs later perhaps due to increased recall of the lack of drug in the prior CPP test experience.

### CA1 5-HT1B Receptors in Memory Retrieval

The role of 5-HT1B receptors in memory is more complicated, and appears to be dependent on the type of memory task and cognitive demand. Mice lacking 5-HT1B receptors show enhanced acquisition of the hippocampus-dependent memory task, Morris water maze, but impaired working memory in the radial arm maze (Buhot et al., 2003). However, 5-HT1B receptor antagonists in CA1 promote memory formation (Cai et al., 2013) and facilitate avoidance learning (Eriksson et al., 2008). Inverse agonists of 5-HT1B receptors, or the 5-HT1B antagonist GR127935, administered post-training can also facilitate learning in associative autoshaping learning tasks (Meneses, 2001). We hypothesize, in our system, that 5-HT1B is acting to enhance drug “memory” recall (e.g. cocaine-seeking on ED1), and blockade of these receptors by GR127935, decreases recall; as shown by decreased cocaine-seeking on ED1.

### CA1 5-HT1A and 5-HT1B Receptors in Drug Memory

Our studies using combined antagonism of 5-HT1A/1B (WAY100,635 plus GR 127935) replicated earlier studies that aimed to more selectively mimic serotonergic actions of S-propranolol (Kohtz and Aston-Jones, 2017). Given the summarized effects of 5-HT1A and 5-HT1B receptors to inhibit consolidation and facilitate recall of drug-associated memories, respectively, administering antagonists of the 5-HT1A and 1B receptors together may prevent both of these actions in tandem. Therefore, on ED1, or during testing of cocaine CPP, decreased recall and cocaine-seeking behavior is observed (due to 5HT1A antagonism). However, the effects of ED1 manipulations on persistence tested 2 weeks later are comparable with the effects of combined antagonism during CPP acquisition, where we observed enhanced memory consolidation. We interpret our results to indicate that 5-HT1A and 5-HT1B receptors in the dHPC are activated on ED1 to decrease consolidation of the absence of drug and enhance contextual recall, respectively, driving increased cocaine-seeking behavior and subsequently maintaining seeking behavior 2 weeks later (Fig.4, Left). When this signal is blocked, either by antagonism of 5-HT1 receptors in the dHPC (Fig.4, Middle) or inhibiting serotonin signaling to the dHPC via DREADDs, we decrease recall of the drug-associated context thereby reducing cocaine-seeking behaviors on ED1, and enhance consolidation of the novel information about the context (e.g. absence of cocaine), thereby producing a persistent effect following 2 weeks of abstinence (Fig.4, Right).

### Conclusions

The present findings indicate that initial abstinence may be a critical time point for targeting subsequent drug-seeking behavior. While these behaviors differ substantially between males and females, the serotonin system may be uniquely involved in both. Thus, targeting drug seeking behavior during initial abstinence via therapeutic interventions that manipulate contextual memory or serotonin signaling has strong treatment implications for both sexes.

## Acknowledgements

We thank members of the Aston-Jones lab for their thoughtful comments on the project. We thank students Belle Lin and David Goldmeier for their technical assistance in completing this work.

## Author Contributions

A.S.K. and G.A.J. designed experiments. A.S.K. performed experiments and data analyses and A.S.K. and G.A.J. developed protocols. A.S.K. and G.A.J. prepared the manuscript. Correspondence and requests for materials should be addressed to G.A.J.

## Figure Legends

**Figure 1. 5-HT1A/1B receptor antagonism in dorsal hippocampal CA1 persistently reduced cocaine-seeking in both males and females. (A)** Timeline of experimental conditions for each rat tested in b-d and g-h. **(B)** Cocaine self-administration of male (0.20 mg/infusion) and female (0.16mg/infusion) rats. Different doses were chosen for males and females to produce similar active lever pressing behaviors. * indicates days in which female lever pressing is significantly greater than males, *p* <0.05. **(C)** Daily cocaine intake by mg/kg body weight of of male (0.20 mg/infusion) and female (0.16mg/infusion) rats. * indicates days in which female overall intake is greater than males, *p* <0.05. **(D)** Administration of WAY/GR to dorsal CA1 10min prior to ED1 testing decreased cocaine-seeking in both males and females on ED1, and persistently decreased seeking when tested 2 weeks later on WD15 (with no additional injections). * indicates significant differences from aCSF administered controls. All CA1 injections were made only prior to ED1. **(E)** Behavioral timeline for testing the effects of WAY/GR on WD1 in the home cage on later cocaine-seeking behavior tested at WD15. **(F)** Administration of WAY/GR to dorsal CA1 on WD1 in the home cage had no effect on subsequent cocaine seeking tested 2 weeks later. **(G)** Survival curves for extinction behavior following ED2 testing depicted in panel D. Although females had significantly less ED2 cocaine-seeking than did males in Panel D, it took longer for saline administered females to meet extinction criterion of <25 lever presses for 3 consecutive days. Intervention with WAY/GR on ED1 significantly reduced extinction resistance in female, but not male, rats. **(H)** Antagonism of 5-HT1A/1B receptors in dorsal CA1 by WAY/GR had no effect on cued reinstatement behavior following extinction. However, there was a significant main effect of sex wherein females had greater cued-reinstatement behavior than did males. * indicates significant differences from males *p* <0.05.

**Figure 2. Chemogenetic inhibition of dorsal raphe signaling in CA1 persistently reduced cocaine-seeking in both males and females. (A)** A midsagittal section of a rat brain illustrating the pAAV8-hSpn-hM4Di(Gi)-mCherry infusion target region (dorsal raphe, DR) and CNO infusion location (dorsal hippocampus, dHPC). Projections from dorsal raphe were transiently inactivated in CA1 by local microinjection of CNO among axon terminals expressing hM4Di receptors. **(B)** Typical mCherry expression (purple) in dorsal raphe, largely contained within dorsal raphe borders. **(C)** Terminal expression of hM4Di by detection of mCherry tag (red). hM4Di fibers were present in dentate gyrus, CA3, and CA1. Cell nuclei are counterstained with DAPI (blue). Panel at right is magnification of area indicated in low-power at left. **(D)** Behavioral timeline for testing performed in panels E and F. **(E)** Administration of CNO (1mM/1.0uL/side) to dorsal CA1 to transiently inactivate dorsal raphe terminals produced persistent effects on cocaine-seeking behaviors in both males and females. * indicates significant differences from aCSF administered controls. **(F)** Transient inactivation of dorsal raphe terminals in CA1 by local CNO had no effect on cued reinstatement behavior following extinction. However, there was a significant main effect of sex wherein females had greater cued-reinstatement behaviors than males. * indicates significant differences from males.

**Figure 3. 5-HT1A/1B receptors modulate acquisition and recall of cocaine conditioned place preference (CPP) memories. (A)** Timeline for testing of administration of WAY100,635 (WAY; 5-HT1A antagonist) and/or GR127935 (GR; 5-HT1B antagonist) during CPP conditioning days (Acquisition) *or* 10 minutes prior to CPP testing (Retrieval). **(B)** CPP memory with WAY/GR given during CPP acquisition days or CPP expression test. There were no significant differences between control administered saline vehicle during acquisition or retrieval (n=4/sex); therefore, these data were combined. WAY/GR given during acquisition increased CPP preference, and WAY/GR given prior to the test session decreased CPP memory. * indicates significant differences from saline administered controls *p* < 0.05. **(C)** CPP performance after WAY100,635 interventions. WAY given on conditioning days enhanced CPP memory. There was no effect of WAY on CPP memory when administered 10 mins prior to the test session. However, when retrieval administered rats tested 24hr later, within-subjects analyses showed a slight conditioned aversion on retest. * indicates significant differences from saline administered controls *p* < 0.05. **(D)** CPP performance after GR127935 interventions. GR given on conditioning days had no effect on CPP. GR administration 10 mins prior to the test session significantly decreased cocaine CPP. When retrieval administered rats were tested 24hr later, within-subjects analyses reveal a recovery of CPP memory. * indicates significant differences from saline administered controls *p* < 0.05.

**Figure 4. Hippocampus CA1 Serotonin Signaling Driving Cocaine Seeking Persistence. (Left Panel)** 5-HT1A and 5-HT1B receptors in the dorsal hippocampus CA1 pyramidal layer are activated on ED1 to decrease consolidation of the absence of drug and enhance contextual recall, respectively, driving increased cocaine-seeking behavior and subsequently maintaining seeking behavior 2 weeks later. **(Middle Panel)** Antagonism of 5-HT1 receptors in the dHPC decreases recall of the drug-associated context thereby reducing cocaine-seeking behaviors on ED1, and enhances consolidation of the novel information about the context (e.g. absence of cocaine), thereby producing a persistent effect following 2 weeks of abstinence. **(Right Panel)** Blockade of 5-HT signaling from the DR via DREADDs decreases signaling at both 5-HT1A and 5-HT1B receptors in the DH, decreasing recall of the drug-associated context thereby reducing cocaine-seeking behaviors on ED1, and enhancing consolidation of the novel information about the context (e.g. absence of cocaine), thereby producing a persistent effect following 2 weeks of abstinence. 5-HT; serotonin, ED; extinction day, WD; withdrawal day, DR; dorsal raphe.

## TO BE PLACED IN SUPPLEMENTAL INFORMATION

### Novel Object Recognition Task (nOR)

We used a clear acrylic chamber (∼40 × 40 × 30 cm) to test nOR performance. Rats were habituated to the box for one day prior to training for 30 minutes. On day 1, rats were exposed to 2 identical objects and allowed to freely explore for 5 mins. Time spent with each object was recorded to determine if rats favored one side of the box. 24 hrs later rats were returned to the box and one object was replaced with a novel object. The percentage of time spent interacting with the novel object indicated object memory performance.

### Effect of 5-HT1A plus 5-HT1B Receptor Antagonist cocktail on Novel Object Recognition Performance

We tested the effects of giving the WAY/GR cocktail into dorsal CA1 on nOR performance to determine if effects of WAY/GR on drug-associated memories can be generalized to other memory types in naïve rats. Administering WAY/GR 10min prior to training in nOR resulted in increased nOR performance (F(1,33)=5.133, *p* <0.05), whereas instead administering WAY/GR 10m prior to testing resulted in decreased nOR performance (F(1,35=25.10, *p* <0.05) of male and female rats. As these effects were small and recapitulated the observed effects in CPP, we did not further investigate the specific role of 5-HT1A and 5-HT1B in nOR.

---

**Supplemental Figure 1. 5-HT1A/1B receptors modulate acquisition and recall of novel object recognition (nOR) memories. (A)** Timeline for testing the effects of WAY100,635 plus GR127935 (5-HT1A and 5-HT1B antagonists, respectively, WAY/GR) administration prior to training (acquisition) or prior to testing (retrieval) on nOR performance. **(B)** nOR performance. There were no significant differences between vehicle controls run in timeline A (n=5/sex) or timeline B (n=5/sex); therefore, these data were combined. WAY/GR given into dorsal CA1 during nOR acquisition enhanced nOR performance, whereas WAY/GR given before the nOR test reduced nOR performance. * indicates significant differences from saline administered controls *p* < 0.05.

